# A functional network signature in the developing cerebellum: evidence from a preclinical model of autism

**DOI:** 10.1101/2021.06.14.448440

**Authors:** María Berenice Soria-Ortiz, Ataúlfo Martínez-Torres, Daniel Reyes-Haro

## Abstract

Autism spectrum disorders (ASD) are pervasive neurodevelopmental conditions detected during childhood when delayed language onset and social deficits are observed. Children diagnosed with ASD frequently display sensorimotor deficits associated with the cerebellum, suggesting a dysfunction of synaptic circuits. Astroglia are part of the tripartite synapses and *postmortem* studies reported an increased expression of the glial fibrillary acidic protein (GFAP) in the cerebellum of ASD patients. Astroglia respond to neuronal activity with calcium transients that propagate to neighboring cells, resulting in a functional network response known as a calcium wave. This form of intercellular signaling is implicated in proliferation, migration, and differentiation of neural precursors. Prenatal exposure to valproate (VPA) is a preclinical model of ASD in which premature migration and excess of apoptosis occur in the internal granular layer (IGL) of the cerebellum during the early postnatal period. In this study we tested calcium wave propagation in the IGL of mice prenatally exposed to VPA. Sensorimotor and social deficits were observed and IGL depolarization evoked a calcium wave with astrocyte recruitment. The calcium wave propagation, initial cell recruitment, and mean amplitude of the calcium transients increased significantly in VPA-exposed mice compared to the control group. Astrocyte recruitment was significantly increased in the VPA model, but the mean amplitude of the calcium transients was unchanged. Western blot and histological studies revealed an increased expression of GFAP and higher astroglial density. We conclude that the functional network of the IGL is remarkably augmented in the preclinical model of autism.

## 1. Introduction

Autism Spectrum Disorder (ASD) refers to a group of neurodevelopmental disorders characterized by social impairment, communication deficits, stereotypies, and repetitive behaviors. ASD is diagnosed by the age of two and the prevalence is about 1% of the global population, but little is known about the neurobiology of the disorder, particularly before diagnosis. Motor disorders associated to the cerebellum are frequently observed in patients diagnosed with ASD before the onset of language or social deficits (Lloyd et al., 2013; Mcphillips et al., 2014; Mosconi et al., 2015). The cerebellum is a site of ASD gene-associated co-expression during early postnatal development, especially in the granular layer (Menashe et al., 2013; Wang, et al., 2014). Granule cells (GCs) of the cerebellum represent nearly half of the neurons of the rodent or primate brain and integrate multiple sensory modalities (Wang, et al., 2014). GC axons give rise to parallel fibers and synapse onto Purkinje cells (PCs). Climbing fibers that project from the inferior olivary nucleus to PCs can drive the plasticity of the resulting signal, since there is a sole inhibitory output from PCs into the deep cerebellar nuclei which in turn project to the thalamus and other brain regions (Huang et al., 2013; Wang et al., 2014). PC loss (Skefos et al., 2014) and the increased expression of astroglial markers, such as aquaporin-4, connexin 43 and GFAP have been reported in *postmortem* studies of ASD patients (Edmonson et al., 2014; Fatemi et al., 2008; Laurence & Fatemi, 2005). Neuron-glia communication is required for normal functioning of the brain during early neurodevelopment and through life. Astroglia are part of the tripartite synapses and respond to neuronal activity with calcium transients that propagate to neighboring cells, a signaling mode known as a calcium wave (Perea & Araque, 2005; Schipke & Kettenmann, 2004). This form of intercellular communication is implicated in the proliferation, migration and differentiation of neural precursor cells (Weissman et al., 2004). Bioinformatic studies have shown that ASD gene-associated co-expression networks are highly expressed in the cerebellar granule layer and that genes involved in calcium signaling form one of the most relevant interaction nodes (Menashe et al., 2013; Zeidán-Chuliá et al., 2013). However, it is unknown whether neuron-glia communication associated with calcium signaling is disturbed in the early neurodevelopment of the autistic brain. Murine models of ASD open the possibility to investigate this problem in a developmental window prior to diagnosis. Prenatal valproate (VPA) exposure is a commonly used preclinical model of ASD to explore mechanistic and therapeutic investigations (Nicolini & Fahnestock, 2018; Varman et al., 2018; Wang et al., 2018). VPA is a mood stabilizer and anticonvulsant drug, but prenatal administration in humans results in linguistic, motor and cognitive deficits (Moore et al., 2000). The VPA model reproduce these deficits including reduced dendritic arborization of PCs, delayed GC precursor migration and neuronal apoptosis (Ingram et al., 2000; Rodier et al., 1997; Varman et al., 2018; Wang et al., 2018). Thus, the aim of this study was to test neuron-glia communication associated with calcium signaling in the IGL of mice prenatally exposed to VPA.

## 2. Experimental Procedures

### 2.1. Experimental animals and ethical standard

Mice were handled according to the National Institute of Health’s Guide for the Care and Use of Laboratory Animals and the Institutional Committee on Animal Care and Use of Laboratory Animals of the Institute of Neurobiology, UNAM. Briefly, CD-1 or GFAP-eGFP transgenic mice (Nolte et al., 2001) were mated and pregnancy was confirmed by a vaginal plug corresponding to embryonic day 0 (E0). The pregnant mice were housed individually under a 12h/12h light/dark cycle with controlled temperature, and food and water *ad libitum.* A single intraperitoneal injection (IP) of sterilized saline solution (0.9%) or VPA (Sigma-Aldrich, St. Louis, MO, USA) was administered at E12.5 (Varman et al., 2018). VPA was dissolved in sterilized saline solution (0.9%) as follows: 500 mg/Kg for CD-1 & 300 mg/Kg for GFAP-eGFP dams. After birth, the pups were weaned at postnatal day 21 (P21) as standard procedure. Only male pups were used for this study based on ASD incidence (4:1) (Kim et al., 2013; Melancia et al., 2018; Perez-Pouchoulen et al., 2016).

### 2.2. Behavioral testing

#### 2.2.1. Latency to reach the nest

The nest-seeking test was carried out at P8 for both CTL and VPA experimental groups (ten litters, n_CTL_ = 35; thirteen litters, n_VPA_ = 47). This test was also performed on three litters from the GFAP-eGFP strain (two males per litter for each experimental group). The pups were placed in the center of a 35 × 20 cm^2^ plastic cage containing clean bedding and home bedding in opposite sides (~5 cm width) with 25 cm of separation. Olfactory cues were prevented by cleaning the cage after each trial. The latency to reach the nest was recorded immediately after the mouse’s head touched the home bedding (Roullet et al., 2010; Schneider & Przewłocki, 2005; Varman et al., 2018).

#### 2.2.2. Righting reflex

The righting reflex is a mouse pup’s motor ability to flip onto its feet from supine position. At P9, the pups were placed on their backs on a flat surface and held in that position for 5 s. Next, they were released and the time it took them to return to prone position was recorded for the CTL (n_CTL_ = 16 from four litters) and the VPA (n_VPA_ = 12 from tree liters) experimental groups (Feather-Schussler & Ferguson, 2016; Kazlauskas et al., 2016).

### 2.3. Western Blot analysis

Cerebella from CD-1 male pups (P8) were dissected from four litters, for both the CTL and VPA groups. Briefly, four sets of cerebella for each experimental group (4 cerebella per set, n = 16) were homogenized in iced-cold glycine lysis buffer (in mM: 200 Glycine, 150 NaCl, 50 EGTA, 50 EDTA, 300 sucrose, pH 9.0) and protease inhibitor (Sigma-Aldrich, St. Louis, MO, USA), followed by protein isolation and quantification with a Bradford assay (Bio-Rad, Hercules, CA, USA) (Bradford, 1976). An equal concentration of protein (10 μg) per lane was resolved in a 10% polyacrylamide gel. The proteins were transferred to PVDF membranes, blocked with 5% nonfat dry milk in Tris-buffered saline (TBS), 0.1% Tween 20 (TBS-T) for 3 h at room temperature. The membranes were incubated overnight at 4°C with the primary antibody goat polyclonal anti-GFAP 1:1000 (Santa Cruz, Dallas TX, USA) or rabbit anti-Actin 1:1000 (Santa Cruz, Dallas TX, USA).

The membranes were rinsed three times (15 min/each) with TBS-T and primary antibodies were detected after incubation (3 h) with either rabbit anti-goat IgG-AP (1:2000) or goat anti-rabbit IgG-AP (1:2000) (Santa Cruz, Dallas, TX, USA). Alkaline phosphatase activity was detected with BCIP/NBT AP-conjugate substrate reaction kit (Bio-Rad, Hercules, CA, USA) after post-secondary washes with TBS-T. The images of the Western blot bands were acquired with the Image based Gel Doc™ EZ Gel Documentation System (Bio-Rad, Hercules, CA, USA). Optical density was calculated with Image Lab 3.0 software (Bio-Rad, Hercules, CA, USA) and normalized with the β-Actin bands (Varman et al., 2018).

### 2.4. Histology

Histological studies were performed in GFAP-eGFP transgenic mice at P8 (n_CTL_ = 3, n_VPA_ = 4, 3 litters each group). Briefly, mice were deeply anesthetized with an IP injection (100 mg/Kg) of pentobarbital and intracardially perfused with chilled (4°C) paraformaldehyde (PFA 4%) in 0.1 M phosphate-buffered saline (PBS, pH 7.4) as previously described (Varman et al., 2018). Brains were carefully isolated and kept in PFA for another 24 h, then washed and cryoprotected in 30% of sucrose at 4°C. Brains were frozen in polyvinyl alcohol-polyethylene glycol medium (Tissue-Plus, Fisher Health Care) at −80°C and horizontal sections (35 μm) including Crus I/II, were obtained with a cryostat (Leica CM1850). Sections were stored in cryoprotectant solution (30% ethylene glycol-30% sucrose in PBS) at −20°C. The histological sections were washed three times (10 min/each) with PBS, counterstained with 4’, 6-diamidino-2-phenylindole (DAPI, 1:16000) and mounted with Vecta-Shield (H1000, Vector Laboratories, Burlingame, CA, USA) (Varman et al., 2018).

### 2.5. Cell counting

Three images per slice from the internal granular layer (IGL) of the Crus I/II regions (n_CTL_ = 3, n_VPA_ = 4, in triplicated samples) were acquired with an ApoTome microscope and a Zeiss LSM 780 confocal microscope (Zeiss, Germany). Images were edited into 50 x 200 μm rectangles using Zen2012 Blue Edition software and processed in ImageJ. Cell somas were counted automatically in CellProfiler 2.1.1 software. Data were analyzed in OriginPro 7.0 and presented as the number of immunoreacted cells per 1 mm2. Statistical analysis was performed with a Shapiro-Wilk for normal population testing and then a Student’s t test for two samples previously tested.

### 2.6. Sulforhodamine B staining of astrocytes

Mice were intraperitoneally injected with sulforhodamine B (SRB; 20 mg/Kg) 4 h prior the brain slice sectioning. This approach was previously reported for astrocyte staining (Appaix et al., 2012; Nimmerjahn et al., 2004). We further corroborated the selectivity of SRB for astrocytes in the GFAP-eGFP transgenic mouse line. Our results showed that 84 ± 2% of GFAP-eGFP^+^ cells incorporated SRB in the IGL of Crus I/II regions (n = 12, N = 3, in triplicated samples). We conclude that SRB stains mainly astrocytes (see suppl. Fig. 1).

### 2.7. Acute brain slice preparation and calcium imaging

Brain slices were obtained as previously described (Labrada-Moncada et al., 2020; Reyes-Haro et al., 2010, 2013). Briefly, acute cerebellar slices were prepared from 7-to 9-day-old (P7-P9) CD-1 or GFAP-eGFP mice (Nolte et al., 2001). After decapitation, the brain was immediately removed and coronal slices (250 μm) containing Crus I/II regions were obtained with a vibratome (VT1000s, Leica) and transferred to ice-cold oxygenated artificial cerebrospinal fluid (aCSF, in mM: 134 NaCl, 2.5 KCl, 2 CaCl_2_, 1.3 MgCl_2_, 26 NaHCO_3_, 1.25 K_2_HPO_4_, 10 glucose, pH = 7.4). Two slices per animal were obtained and stored at room temperature in oxygenated aCSF for at least 30 min, followed by incubation with the Ca^2+^ indicator dye Fluo-4 AM (10 μM, AAT Bioquest, Sunnyvale, CA, USA) for another 30-45 min at 37°C. The slices were washed with aCSF for 30 min, transferred to the recording chamber and perfused with oxygenated aCSF (2 ml/min) at room temperature (20-22°C). Calcium imaging experiments were performed under a cooled camera (SensiCam; PCO.Edge 4.2, Kelheim, Germany) coupled to an Olympus upright microscope (BX51WI, Miami, FL, USA) and a LED module (X-Cite X-LED1 lumen Dynamics Fremont, CA, USA; BDX: 450-495 nm for Fluo-4AM and GYX: 540-600 nm for SRB). The calcium waves were evoked by depolarization (20 pulses of 200 μA at 10 Hz evoked with a DS3 Isolated Current Stimulator, Digitimer Ltd, Fort Lauderdale, FL, USA) with a pipette pulled from thin-walled borosilicate glass (outer diameter 1.5 mm, inner diameter 0.87 mm) with a P97 puller (Sutter Instruments, Novato, CA, USA). The stimulation electrode (with a tip opening of ~20 μm) was filled with aCSF and placed on top of the slice, gently touching the upper cells of the IGL. Then, the slice was allowed to recover from mechanical stress for at least 5 min. The image acquisition protocol consisted of 200 s at 1 Hz. The first 15 s were defined as the pre-stimulus window before depolarization. Image analyses and processing were performed with ImageJ/FIJI software. The cells recruited within the calcium wave were visualized after subtraction of pixel values from the previous image. All the recruited cells were displayed after adding up all the images with the corresponding background subtraction (Haas et al., 2006; Labrada-Moncada et al., 2020; Reyes-Haro et al., 2010). The functional extension (maximum length) and number of recruited cells within the calcium wave were estimated using concentric rings with 50 μm increments around the site of stimulation. The speed of propagation was defined as (maximum length) / (t_1_-t_0_); where the t0 was the time when the first cells were activated immediately after the stimulus and t_1_ was the time when the farthest cells were activated. The threshold of the evoked intracellular calcium transients ([Ca^2+^]i) was estimated as *ΔF* = *ΔF*/*F*_*b*_, where *ΔF* is the relative change of the fluorescence over the basal fluorescence *F*_*b*_. The average of the 15 frames at the beginning of the recording was defined as F_b_. The area under the curve was estimated with the trapezoidal rule, a numerical integration method that considers several small trapezoids under the curve. The arbitrary units represent the amplitude of the response ([Ca^2+^]i) through a time scale (in seconds).

### 2.8. Statistical analysis

Statistical analysis was performed using Origin Lab 8.0 software (Origin Laboratories, Northampton, MA, USA) and includes the Shapiro-Wilk test to determine a normal distribution, t-Student or Mann-Whitney tests to analyze differences between CTL and VPA groups in populations with normal and non-normal distributions. Data are reported as mean ± SEM. P values < 0.05 were considered significant.

## 3. Results

### 3.1. Sensorimotor and social skills are deteriorated by prenatal exposure to VPA

The VPA model of autism reproduces behavioral symptoms associated to the disorder. In this work, we tested sensorimotor and social skills through postnatal development of CTL and VPA mice. Our data showed sensorimotor and social skills deficits associated to a prenatal exposure to VPA during postnatal neurodevelopment (Fig. 1).

**Figure 1.**
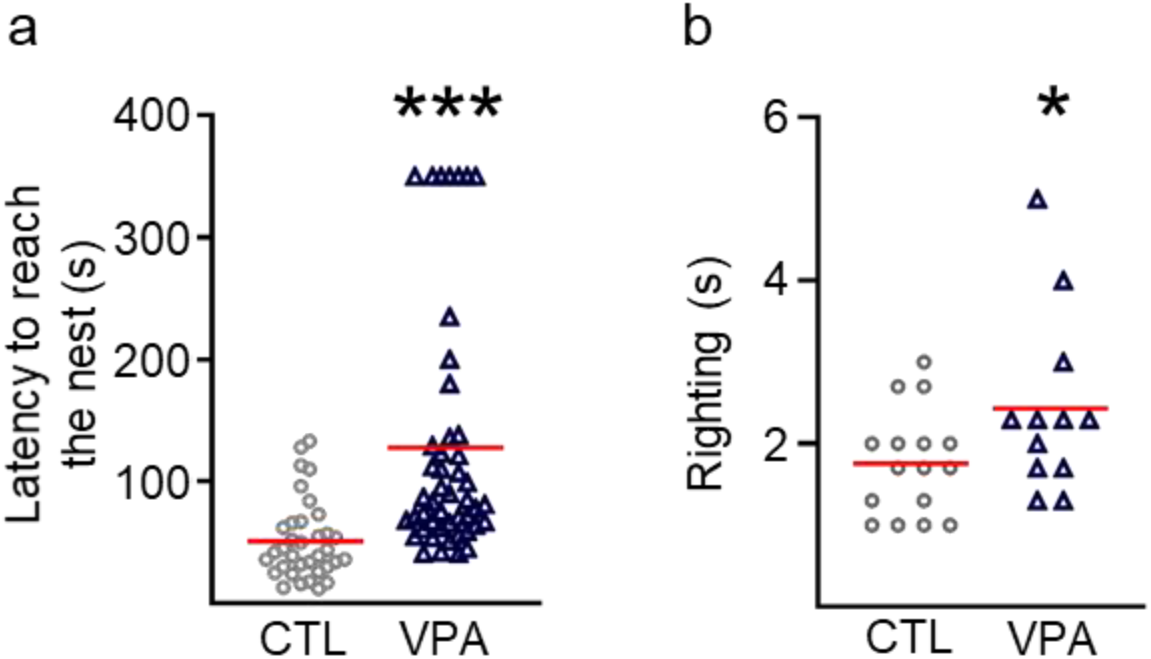
Sensorimotor and social deficits in the VPA model of autism. Graphs show the latency to reach the nest (a) performed at P8 (n_CTL_= 35, n_VPA_= 47, N_VPA_ = 13) and the righting reflex latency (b) performed at P9 (n_CTL_= 16, n_VPA_= 12). Control (CTL); valproate (VPA). Data analyzed by Shapiro–Wilk, Mann–Whitney U, and Student’s t-test, *p < 0.05, **p < 0.001, ***p < 0.0001. Values are mean ± SEM.

#### Latency to find the nest

The latency to reach the nest was increased in VPA-treated pups (127.7 ± 14.9 s, n_VPA_= 47, N_VPA_ = 13, p = 3E-7) when compared to the CTL group (51.2 ± 5.5 s, n_CTL_= 35, N_CTL_ = 10; Fig. 1a).

#### Righting reflex

The righting reflex latency was augmented in VPA-treated pups only at P9 (2.43 ± 0.3 s, n_VPA_= 12, N_VPA_ = 3, p = 0.05) when compared to CTL group (1.75 ± 0.16 s, n_CTL_= 16, N_CTL_ = 4; Fig. 1b).

### 3.2. The functional network of IGL is increased in the VPA model

Once we determined that VPA-treated mice showed autistic-like deficits, we locally evoked a calcium wave depolarizing *in vivo* slices of the IGL of the Crus I / Crus II regions of the cerebellum to analyze if the maximum length of propagation, determined by the number of Fluo-4AM recruited cells was altered in the VPA-model (Fig. 2a). Data showed no significant differences of the total number of recruited cells between both experimental groups (data not showed). Nevertheless, the maximum length of the calcium wave propagation was increased (+29%) in VPA-treated mice (382 ± 22 μm, n_VPA_ = 10, N_VPA_ = 9, p = 0.01) when compared to the CTL group (295.9 ± 24, n_CTL_ = 10, N_CTL_ = 7) (Fig. 2c), suggesting a different recruited cell distribution in the VPA-model.

**Figure 2.**
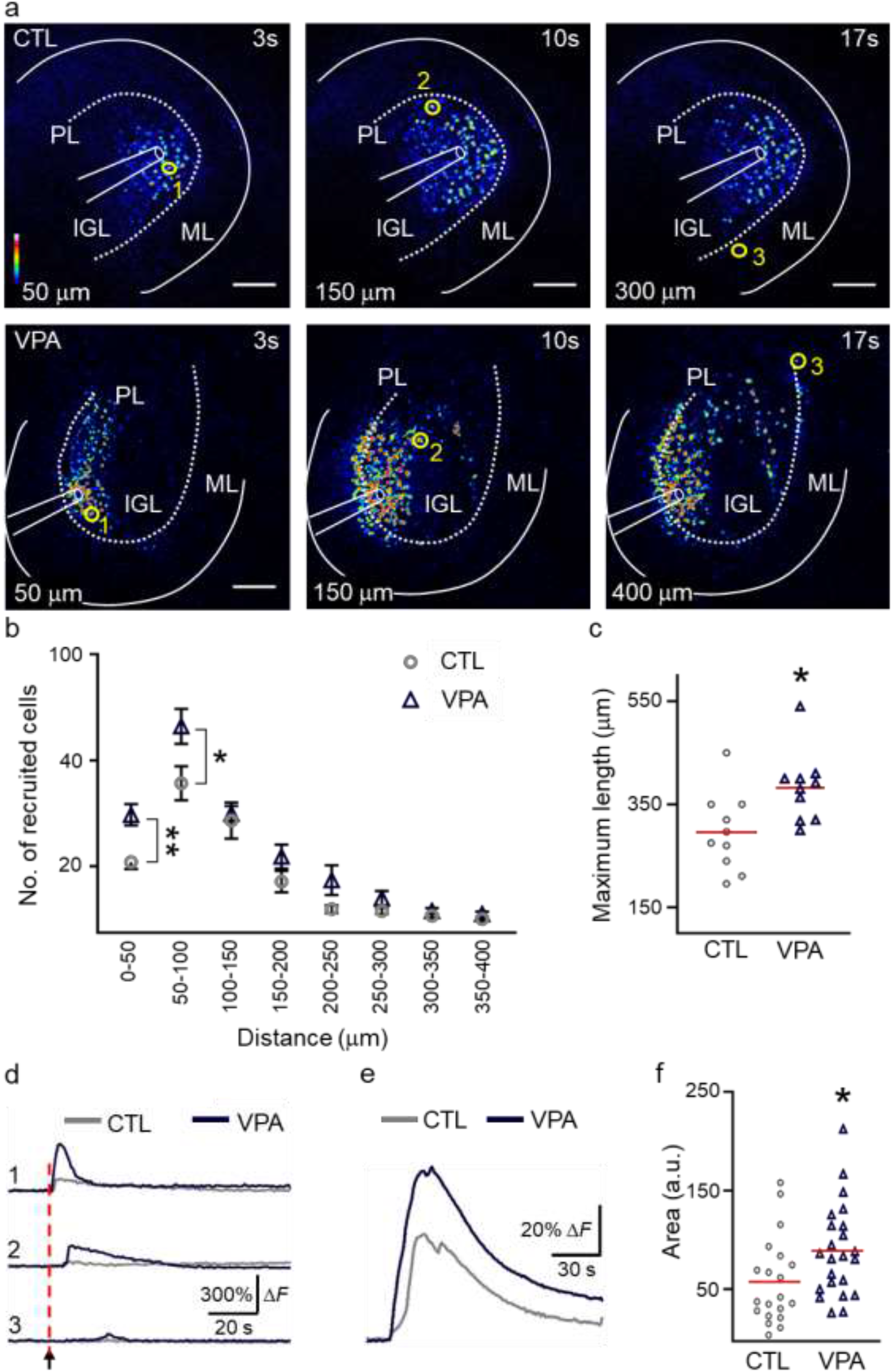
The calcium wave is augmented in the VPA model. (a) Evoked calcium waves in control (CTL) and VPA groups. Pseudocolors (from blue to white) represent the intensity of the fluorescent calcium indicator (Fluo4-AM). Scale bar 100 μm. (b,c) Graph of the recruited cells per distance (n_CTL_= 10, N_CTL_= 7, n_VPA_= 10, N_VPA_= 9). (c) Maximum length of the calcium wave. (d) Example of three cells recruited at different distances in the evoked calcium wave (yellow circles: cell 1, 50 μm; cell 2, 150 μm; cell 3, 300 or 400 μm). (e) The mean amplitude of the calcium transient was estimated from all the recruited cells within the calcium wave in both CTL (CellsCTL = 1222, n_CTL_= 22, N_CTL_= 13) and VPA (Cells_VPA_ = 967, n_VPA_= 23, N_VPA_= 9) experimental groups. (f) Summary of the mean amplitude ot the calcium transient obtained from all the recruited cells from CTL and VPA groups. IGL, internal granular layer; PL, Purkinje layer; ML, molecular layer. Data analyzed by Shapiro–Wilk, Mann–Whitney U, followed by two-way ANOVA, *p < 0.05. Values are mean ± SEM.

Interestingly, our analysis showed that the number of recruited cells increased significantly (+82%) within the initial distance; 0-50 μm (CTL: 22 ± 3 cells, VPA: 40 ± 4 cells, p < 0.001). The second distance (50-100 μm) showed a similar increase (+43 %; CTL: 51 ± 6 cells, VPA: 73 ± 7 cells, p < 0.02), while no significant differences were found for the remaining distances (Fig. 2b and suppl. table 1).

The next step was estimating the mean amplitude of the evoked calcium transients for both experimental groups. Data showed that the largest amplitudes were recorded from the cells located within the initial radius (0-100 μm) and the amplitude of the calcium transients decreased proportionally to the distance from the origin of the depolarization (Fig. 2d). Besides, the mean amplitude of the calcium transients increased 53% (Cells_CTL_ = 1222, n_CTL_= 22, N_CTL_= 13, Cells_VPA_ = 967, n_VPA_= 23, N_VPA_= 9) for the VPA group (89 ± 10 *a. u.*, p = 0.01) when compared to the CTL group (58 ± 10 *a. u.*) (Fig. 2e, f).

Based on these results we conclude that the calcium wave reached longer distances, recruited more cells and the mean amplitude of the calcium transients was increased in the VPA model.

### 3.4. Astrocyte recruitment within the calcium wave is augmented in the VPA model

The IGL is highly populated by granular cells and velate astrocytes, which can also be activated by depolarization. We used SRB, a fluorescent dye that is preferentially incorporated by astrocytes (>80% GFAP^+^ cells, n = 8, N = 3) and was observed that the evoked calcium wave recruited SRB^+^ cells (>50%; n = 8, N = 5) (Suppl. Fig. 1). These results indicate that approximately half of the cells recruited by the evoked calcium wave were astrocytes. Our next step was to test astrocyte recruitment and the corresponding mean amplitude of the evoked calcium transient in the VPA model. Our results showed that SRB^+^ cell recruitment was significantly augmented (+168%) within 100-150 μm ratio (CTL: 8.2 ± 1.6 SRB^+^ cells, VPA: 22 ± 6.5 cells, p < 0.03); but the mean amplitude of the evoked calcium transient was 1671 ± 578 a. u. and 1629 ± 6.336 a. u., for the CTL (Cells_CTL_ = 299, n_CTL_= 6, N_CTL_= 4) and VPA (Cells_VPA_ = 340, n_VPA_= 4, N_VPA_= 3), respectively ( p < 0.4) (Fig. 3 and suppl. table 2). Based on these results we conclude that the evoked calcium wave augmented astrocyte recruitment without changes in the mean amplitude of the calcium transient, in the VPA model.

**Figure 3.**
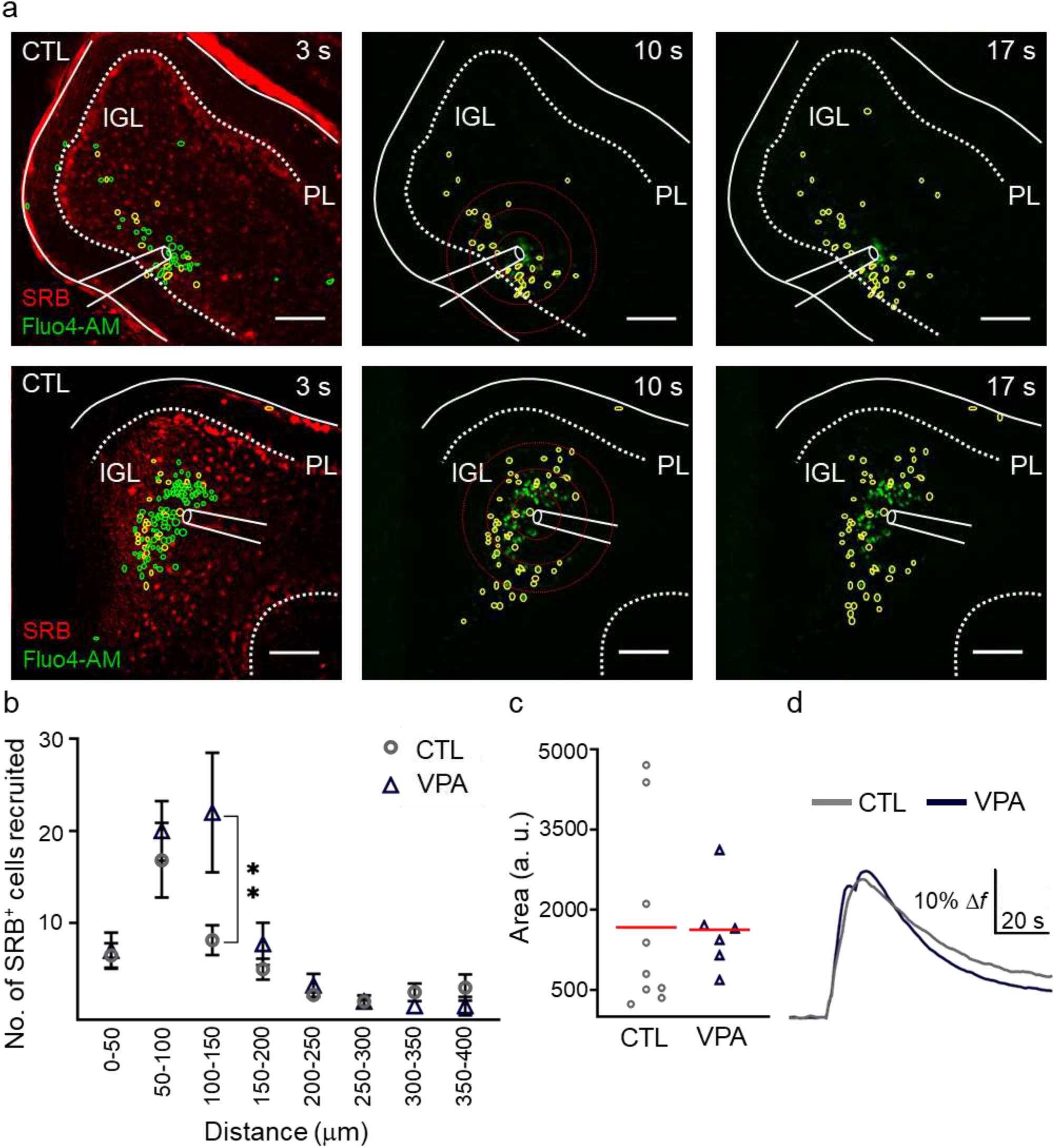
Astrocyte recruitment is augmented within the calcium wave in the VPA model. (a) Representative CTL and VPA images of an evoked calcium wave (Fluo4-AM in green) in IGL; at three different times (3, 10, and 17 seconds). Fraction of Fluo-4AM cells incorporated SRB (yellow rois), the rest of the recruited cells (green rois). Scale bar 100 μm. (b) Histogram of SRB^+^ cell recruitment per distance (n_CTL_ = 6, n_VPA_ = 4, and N_CTL_ = 3 and N_VPA_ = 3). (c) The mean amplitude of the calcium transient was estimated from all SRB+ recruited cells within the calcium wave in both CTL (Cells_CTL_ = 299, n_CTL_= 6, N_CTL_= 4) and VPA (Cells_VPA_ = 340, n_VPA_= 4, N_VPA_= 3) experimental groups. (d) Summary of the mean amplitude of the calcium transient obtained from all the recruited cells from CTL and VPA groups. IGL, internal granular layer; PL, Purkinje layer. Data analyzed by Shapiro–Wilk, and Student’s t-test, **p < 0.05. Values are mean ± SEM.

### 3.5. GFAP expression and astroglial density are increased in the VPA model

The expression of GFAP was tested by Western blot with protein samples isolated from cerebella from both experimental groups (CTL and VPA) in 4 independent sets of 4 cerebella each (n_CTL_= 16, n_VPA_ = 16, each group). Our results showed that the expression of GFAP was increased in the VPA model (28%) compared to CTL group (Fig. 4a). Our next step was to test if higher expression of GFAP correlates with a rise in the density of GFAP^+^ cells. Thus, GFAP^+^ cells were counted in the posterior regions of the cerebellum of GFAP-eGFP transgenic mice, Crus I, Crus II and the vermis region of the VII lobule. The density of GFAP^+^ cells increased 255% in the VPA model (1773 ± 251 cells/mm^2^, n = 4, p = 0.005) when compared to CTL (500 ± 102 cells/mm^2^, n = 3) (Fig. 4b, c). We conclude that GFAP expression and astrocyte density are increased in the VPA model.

**Figure 4.**
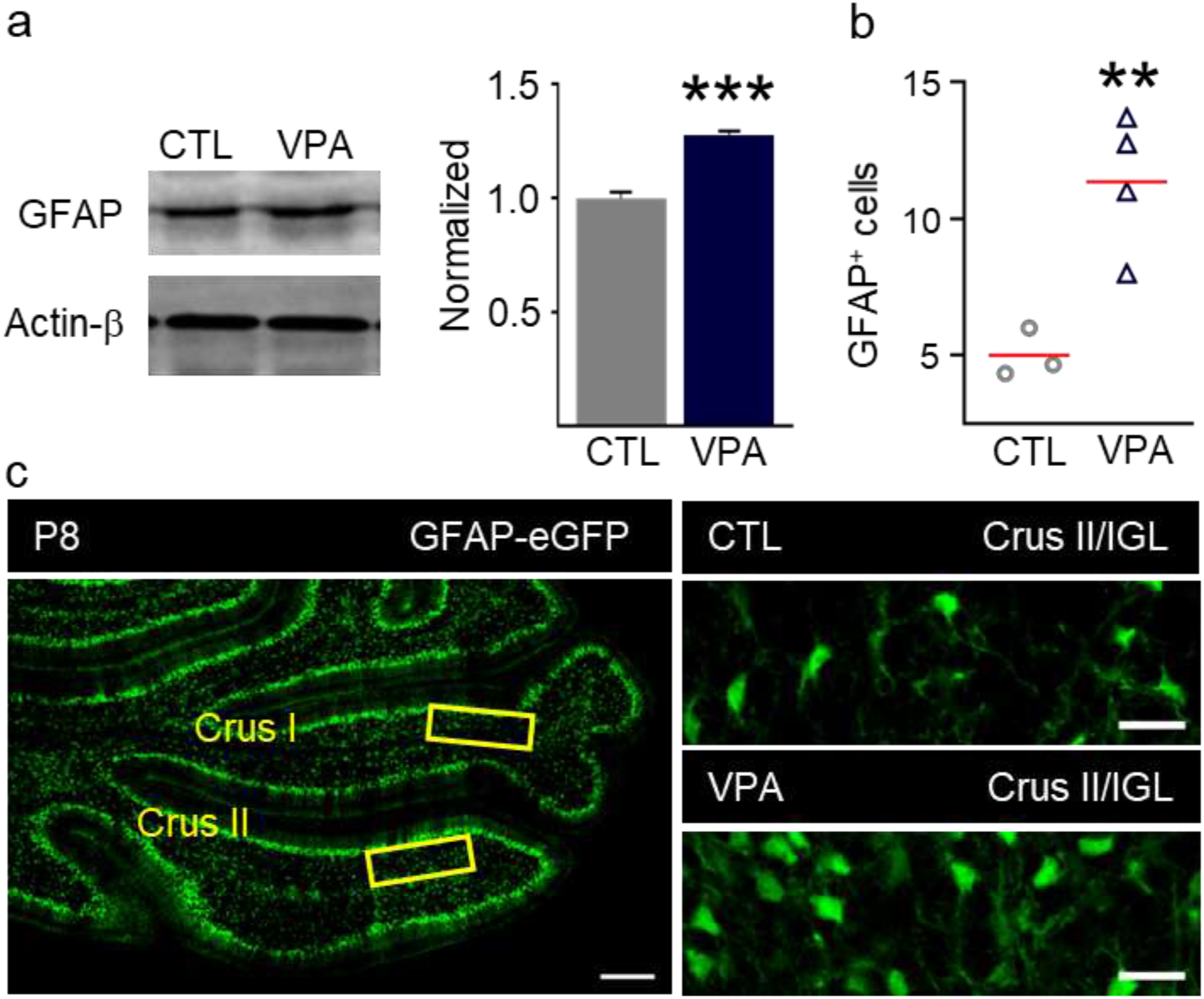
GFAP expression and astrocyte density in the IGL are increased in the preclinical model of autism. (a) Representative Western blot showing the expression of GFAP protein (50 KDa) for CTL and VPA experimental groups. Actin-β (43 KDa) was used as internal control. The band intensities were normalized for CTL (n = 16) and VPA (n = 16) experimental groups (b,c). The density of GFAP+ cells is increased in the VPA model. Cell counting was performed in the cerebellar regions Crus I and Crus II of GFAP-eGFP transgenic mice (n_CTL_= 3, n_VPA_= 4). (c) Left, the horizontal section from the cerebellum shows Crus I and Crus II regions and the internal granular layer (IGL, yellow rectangles). Scale bar 100 μm. Right, representative sections of IGL in Crus II showing GFAP+ cell density CTL and VPA experimental groups. Scale bar 20 μm. Data analyzed by Shapiro–Wilk and Student’s t-test, **p < 0.01, ***p < 0.0001. Values are mean ± SEM.

## 4. Discussion

Brain plasticity resides in the capabilities of nerve cells to generate and convey signals. Glial cells sense and respond to neuronal activity with intracellular calcium transients that can propagate to neighbor cells in a wave-like fashion (Deitmer et al., 2006; Scemes & Giaume, 2006). In this study we tested the cellular response of the IGL to depolarization in a preclinical model of autism, particularly the propagation and velocity of the evoked calcium wave, the number of recruited cells and the mean amplitude of the evoked calcium transients. These functional analyses were performed on mice that showed sensorimotor and social deficits, as well as increased expression of GFAP and astroglial density in the cerebellum due to prenatal exposure to VPA. The cell recruitment, the calcium-wave propagation, and the mean amplitude of the recorded calcium transients evoked by depolarization of the IGL were also increased.

### 4.1. Sensorimotor and social deficits

Sensorimotor and social deficits were observed in mice prenatally exposed to VPA. The increased latencies in righting reflex and nest seeking are in agreement with previous studies using the VPA model (Favre et al., 2013; Kazlauskas et al., 2016; Roullet et al., 2010; Schneider & Przewłocki, 2005; Tartaglione et al., 2019; Varman et al., 2018; Wang et al., 2018). In those studies, autistic subjects were consistently identified based on these neurodevelopmental delays. The sensorimotor delays in the VPA model of ASD correlate with motor deficits observed in ASD children prior to the onset of social or verbal disorders (Lloyd et al., 2013). Experimental data support the examination of early motor deficits as a potential indicator of ASD (Brisson et al., 2012; Sacrey et al., 2015).

### 4.2. The functional network of the IGL

Abnormal genesis and development of neural networks are linked to ASD. Neuron-glia communication is required for normal functioning of the brain during early neurodevelopment and throughout life. The intracellular Ca^2+^ transients occur spontaneously or in response to neuronal depolarization, and propagation to neighboring cells results in a calcium wave (Apuschkin et al., 2013; Hoogland et al., 2009; Kumada & Komuro, 2004; Verkhratsky et al., 2012). Thus, we evoked a calcium-wave by depolarizing the IGL and determined the number of recruited cells. Our results showed that >50% of the recruited cells were astrocytes, suggesting that the other half correspond to GCs and/or neuronal precursors. In agreement, previous studies showed that calcium waves are ATP-driven and expand passing through glial and GCs of the cerebellum (Apuschkin et al., 2013; Hoogland et al., 2009). Calcium waves are thought to ensure correct wiring of the cerebellar circuits, and the intracellular calcium transients recorded in granular cell precursors are known to be involved with migration through the cerebellar cortex (Apuschkin et al., 2013; Hoogland et al., 2009; Kumada & Komuro, 2004). Regarding the dynamics of the calcium wave, our results showed that the propagation, the cell recruitment, and the mean amplitude of the recorded calcium transients were significantly increased in the VPA model. These results correlate with bioinformatic studies in the ASD context where calcium signaling is the most relevant node in the cerebellum (Zeidán-Chuliá et al., 2013) and ASD-gene associated co-expression networks have the strongest correlation in the cerebellar IGL (Menashe et al., 2013). This is highly relevant considering that GCs represent approximately half of the neurons of the human brain (Wang, et al., 2014). GCs receive excitatory synaptic input from mossy fibers and integrate many different sensory receptive fields. Then, the information is distributed across PCs. The axons of GCs originate the parallel fibers, a glutamatergic input that projects into the molecular layer (ML) and contact the dendrites of PCs. Parallel fiber synapses begin to appear at P7 in mice (Zwaigenbaum et al., 2013) and normal calcium signaling in the IGL is necessary for the correct wiring of the cerebellum (Apuschkin et al., 2013; Hoogland et al., 2009; Kumada & Komuro, 2004). Our results show that the functional network of the IGL is augmented in the VPA model, and the increased calcium signaling observed in this study may correlate with the excessive apoptosis of the GC precursors, previously reported in this preclinical model of autism (Wang et al., 2018). Parallel fibers are the presynaptic input of PCs and abnormal function of IGL network might explain the impaired dendritic arborizations and synaptic transmission observed in PCs (Wang et al., 2018). Dysfunction of the cerebellar circuitry is involved, in part, with the sensorimotor deficits observed in autism during early postnatal development (Wang, et al., 2014).

### 4.3. Astroglial is part of the IGL network

Astroglia organizes the architecture of neural networks, nurture synapses and modulate synaptic activity, promoting their development and maturation in the brain (Auld & Robitaille, 2003; Steinmetz et al., 2006; Verkhratsky et al., 2012). The IGL contains cerebellar glomeruli consisting of granule cell dendrites, Golgi cell axon terminals and mossy fibers, all of them wrapped by velate astrocytes. Granule cells represent about 50% of the neuronal population of the brain, consequently, velate astrocytes represent a major astroglial population within the brain. However, little is known about astroglial integration into the functional network of the IGL. In this study we depolarized the IGL and evoked a calcium wave where half of the recruited cells corresponded to astroglia. This network was augmented in the VPA model with a significant increase in astrocyte recruitment. Accordingly, western blot and histological studies showed an increased expression of GFAP that correlated with an augmented density of astroglia in the IGL of VPA exposed mice. However, previous studies using the VPA model reported no differences in GFAP-protein expression, but a decreased density of astrocytes was observed in older mice (P35) (Bronzuoli et al., 2018; Kazlauskas et al., 2016). A possible explanation for these discrepancies may be that we used sensorimotor deficits as a prognostic tool to identify autistic individuals which were selected for histological and functional analyses. In support of our results, Western blot studies showed increased expression of GFAP, whereas immunofluorescence studies reported increased density of astrocytes in the cerebellum of *postmortem* brains from ASD patients (Edmonson et al., 2014; Laurence & Fatemi, 2005; Vargas et al., 2005). On the other hand, the mean amplitude of the calcium transient was unchanged in astrocytes. However, homeostatic functions of astroglia include calcium buffering of the glomeruli (Coenen et al., 2001) and this task may be disturbed by increased density of this cell type, resulting in an increased calcium transient amplitude in the VPA model. Augmented calcium signaling is known to delay migration of GC precursors and increased apoptosis of GCs is observed in the VPA model (Apuschkin et al., 2013; Kumada & Komuro, 2004; Wang et al., 2018), consequently, a reduced dendritic arborization of PCs was reported in the VPA model (Wang et al., 2018).

Overall, we conclude that the functional network of the IGL is significantly augmented in the VPA model of autism, which may correlate with neurodevelopmental delays observed in ASD.

## Credit authorship contribution statement

María Berenice Soria-Ortiz: Conceptualization, Methodology, Validation, Formal Analysis, Investigation, Visualization, Writing original draft. Ataúlfo Martínez Torres: Resources, Writing - review & editing. Daniel Reyes-Haro: Conceptualization, Methodology, Supervision, Resources, Funding acquisition, Writing – review & editing.

## Acknowledgments

M. Berenice Soria-Ortiz is a doctoral student from Programa de Doctorado en Ciencias Biomédicas, Universidad Nacional Autónoma de México (PDCB-UNAM) and received a fellowship from Consejo Nacional de Ciencia y Tecnología (CONACYT, 175835). This research was supported by PAPIIT-UNAM grants to Dr. A. Martínez-Torres (IN204520) and Dr. D. Reyes-Haro (IN205718 and IN209121). The authors declare no conflict of interest. We thank the technical support of E. N. Hernández-Ríos, M.L. Lara-Ayala, A. Castilla, L. Casanova, N. Aranda, E. Espino, M. García-Servín & D. Ragu-Varman. We thank LCC Jessica González Norris for editing and proofreading the manuscript. We are indebted to R. Arellano for allowing us to use his histological facilities.

## Figure legends

Suppl. Fig. 1. Sulforhodamine B (SRB) stains GFP^+^ cells from GFAP-eGFP transgenic mouse. (a) SRB signal in Crus II showing the internal granular layer (IGL, red rectangle). Scale bar 100 μm. (b) Zoom of IGL showing SRB^+^ and GFAP^+^ cells. Merge of both images shows cellular overlay (white). Scale bar 20 μm. (c) Summary of the experiments shows that ~80% of GFAP^+^ cells are SRB^+^ (n = 8, N = 3). (d) Schematic representation of a cerebellar coronal slice showing depolarization of IGL. (e) The calcium wave (Fluo4-AM, green) recruited SRB^+^ cells (overlay in yellow). (f) Summary of the experiments shows that > 50% of recruited cells are SRB^+^ (n = 8, N = 5). Scale bar 100 μm. Stim, stimulus; PL, Purkinje layer; WM, white matter. Statistical tests used were Shapiro–Wilk and Mann–Whitney U.

