## Supplemental Table 1 for "A functional network signature in the developing cerebellum: evidence from a preclinical model of autism"

**Suppl. Table 1**

Table 1. Counting of cells per radius

| Radius (μm) | CTL (No. of cells) | VPA (No. of cells) | *p*-value |
|---|---|---|---|
| 0-50 | 21.6 ± 2.6 | 39.5 ± 4 | **0.001**** |
| 50-100 | 51.4 ± 6.4 | 72.9 ± 6.6 | **0.02*** |
| 100-150 | 37.3 ± 6.8 | 39.9 ± 2.9 | 0.7 |
| 150-200 | 14.3 ± 4 | 23.7 ± 4.5 | 0.14 |
| 200-250 | 3.9 ± 1.5 | 14.8 ± 5.6 | 0.07 |
| 250-300 | 3.2 ± 1.2 | 8 ± 2.7 | 0.19 |
| 300-350 | 1.3 ± 0.7 | 3.2 ± 1.1 | 0.14 |
| 350-400 | 0.1 ± 0.1 | 2 ± 0.9 | **0.03*** |
| 400-450 | 0 | 0.2 ± 0.1 | 0.16 |
| 450-500 | 0 | 0.1 ± 0.1 | 0.3 |

Data are presented as mean ± SEM. Sample size was $n_{CTL}$= 10, $N_{CTL}$= 7, $n_{VPA}$= 10, $N_{VPA}$= 9. *p < 0.05, **p < 0.001.
