## Supplemental Table 2 for "A functional network signature in the developing cerebellum: evidence from a preclinical model of autism"

**Suppl. Table 2**

Table 2. Counting of $SRB^+$ cells per radius

| Radius (μm) | CTL (No. of cells) | VPA (No. of cells) | *p*-value |
|---|---|---|---|
| 0-50 | 6.5 ± 1.3 | 7 ± 2 | 0.83 |
| 50-100 | 16.8 ± 4.1 | 20 ± 3.2 | 0.59 |
| 100-150 | 8.2 ± 1.6 | 22 ± 6.5 | **0.03*** |
| 150-200 | 5 ± 1.1 | 7.8 ± 2.3 | 0.29 |
| 200-250 | 2.2 ± 0.4 | 3.3 ± 1.3 | 0.4 |
| 250-300 | 1.5 ± 0.6 | 1.5 ± 0.6 | > 0.9 |
| 300-350 | 2.5 ± 1 | 1 ± 0 | 0.24 |
| 350-400 | 3 ± 1.4 | 1 ± 1 | 0.41 |

Data are presented as mean ± SEM. Sample size was $n_{CTL} = 6$, $n_{VPA} = 4$, and $N_{CTL} = 4$, and $N_{VPA} = 3$. Statistical analysis used was Mann-Whitney. *$p < 0.05$.
