## Supplementary figures and images for "A functional network signature in the developing cerebellum: evidence from a preclinical model of autism"

### Supplemental Figure 1

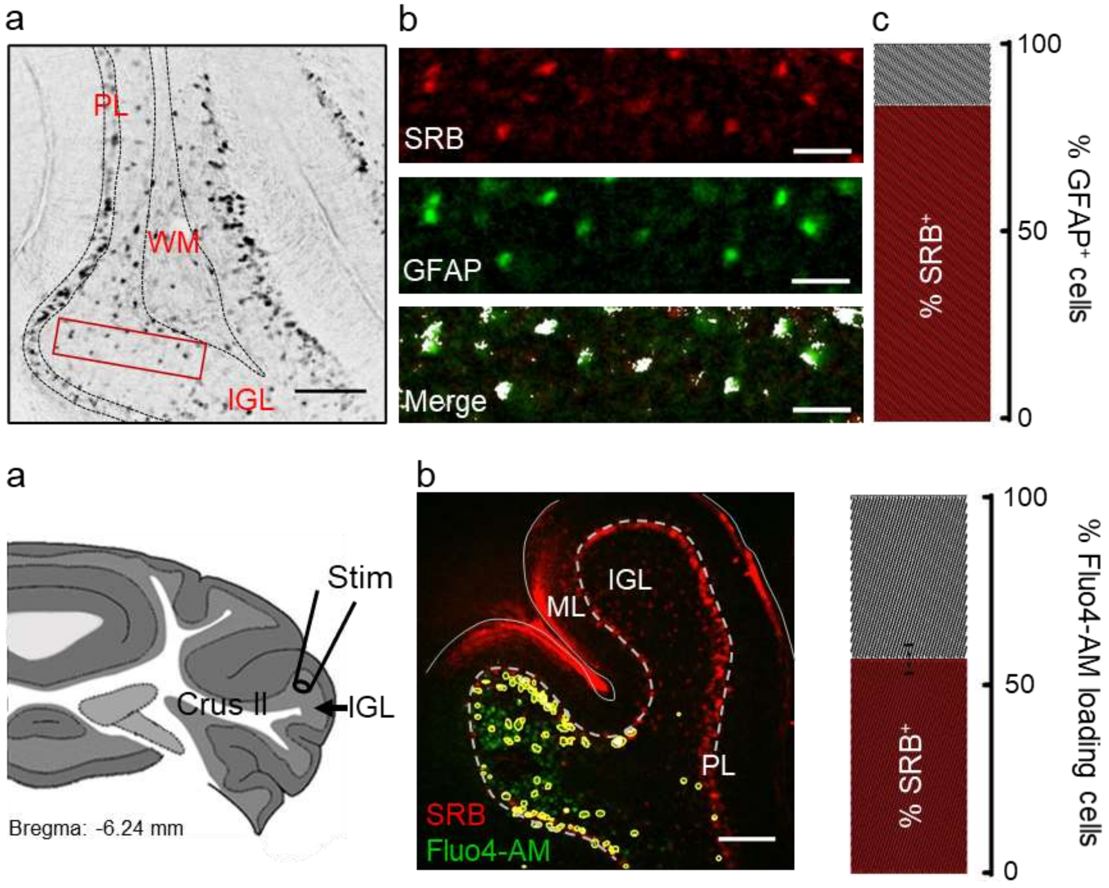
